# Estimating the Designability of Protein Structures

**DOI:** 10.1101/2021.11.03.467111

**Authors:** Feng Pan, Yuan Zhang, Xiuwen Liu, Jinfeng Zhang

## Abstract

The total number of amino acid sequences that can fold to a target protein structure, known as “designability”, is a fundamental property of proteins that contributes to their structure and function robustness. The highly designable structures always have higher thermodynamic stability, mutational stability, fast folding, regular secondary structures, and tertiary symmetries. Although it has been studied on lattice models for very short chains by exhaustive enumeration, it remains a challenge to estimate the designable quantitatively for real proteins. In this study, we designed a new deep neural network model that samples protein sequences given a backbone structure using sequential Monte Carlo method. The sampled sequences with proper weights were used to estimate the designability of several real proteins. The designed sequences were also tested using the latest AlphaFold2 and RoseTTAFold to confirm their foldabilities. We report this as the first study to estimate the designability of real proteins.

## Introduction

Natural protein sequences and structures belong to distinct classes among all potential sequences and structures. A natural protein sequence has a unique global minimum of free energy as its native form, separated in energy from other misfolded states [1]. However, the random amino acid sequences usually don’t have such a feature. In general, protein structures have impressive geometric regularities [2,3], characterized by favored secondary structures and motifs [4] and tertiary symmetry. It has been noted that a small number of folds [5,6] or superfolds account for a huge number of proteins [7]. So, natural proteins are known to fold to a limited number of folds. Some folds are frequently occurring, and lots of sequences can assume that fold. If the sequence capacity of a protein system is big, its structure or function is likely to be resistant to mutation from an evolutionary perspective [17]. Higher sequence capacity has been related to a faster rate of protein evolution [18] and thermophilic adaptation.

This problem has been investigated from the view of “designability” [8-11], which is measured by the number of sequences that the structure can accommodate without loss of thermodynamic stability. Designability has been suggested as an important attribute that contributes to proteins’ functional robustness [8]. Because protein structures are structured in a hierarchy [6,12-14] with four levels ranging from folds to superfamilies, families, and sequences, a protein fold’s designability is determined by the number of families that adopt the fold as their native structure. Many disease-related proteins have folds with few families, and many of these proteins are related to diseases that frequently occur [15]. The size of gene families has also been demonstrated to be substantially linked with designability [16]. However, simply using the number of families under each fold is not a reliable measure for the fold’s designability and its functional robustness because it has been found that the age of a fold correlates with its “usage” among natural proteins. For example, eukaryotic folds found only in human, mouse, and yeast contain approximately an average of 2.5 families, compared to an average of 13.8 families per fold for all human proteins[15]. This observation has two propositions: firstly, since the number of sequences that exist in nature fold to a particular protein structure is not always a good indicator of its designability, we need to estimate the designability of a protein structure to have a better understanding of its functional robustness; second, since new folds have been much less “explored” by nature, there must exist new families, not related to any families found previously, that can be categorized to one of the newer folds. These new protein families may be hosts for some interesting, new functions. It is now possible to design new sequences using a program such as ProDCoNN to target uncharted regions in the foldable sequence spaces specifically.

Many studies on structure designability using lattice model were carried out [10,11, 19-23] where the structures are defined on a 2D or 3D lattice and only two types of residues, hydrophobic (H) and polar (P), are used. Another lattice model was brought by Li et al. using all 20 amino acids with empirically determined interaction potentials (Miyazawa-Jernigan Matrix) [24]. On the other hand, some off-lattice models were proposed [25-28]. Leelananda et al. used small real protein structures from the PDB and employed interaction network representations for those structures to predict their designability [29]. In protein design, using a reduced alphabet would considerably accelerate the search of the sequence space for suitable folders. However, understanding the smallest alphabet necessary for accurate protein design is still an open question [30]. The lattice model result was known to depend on the details of the model. The highly designable lattice structures with a two-letter “amino acid” alphabet are not especially designable with a higher-letter alphabet [31]. Besides, those studies using lattice models or highly coarse-grained energy functions could not quantitatively address the designable space of real protein folds. Therefore atomic design methods are potentially necessary.

In recent years, deep learning methods based on neural networks have dramatically impacted the computational biophysics field, which helps to solve computational protein design (CPD) problems [32-37]. Most recently, Zhang et al. developed ProDCoNN based on a convolutional neural network and reached an accuracy of 42.2% for the test dataset (30% sequence similarity) [38]. Later, Qi et al. designed DenseCPD using DenseNet architecture and improved accuracy to 50.96% [39]. However, there are still no reports about a method of estimating designability using deep neural networks.

In the past, it was always a challenge to testify the eligibility of designed sequences on whether they can fold to a near-native structure because it is impractical to do experimental protein synthesis for thousands of designed sequences. In 2020, AlphaFold2 was developed and achieved remarkable performance in CASP14[40], which is recognized to have solved the protein structure prediction problem at a similar level with experiments. Later RoseTTAFold was also developed, incorporating related ideas, and approach similar accuracy with AlphaFold2[41]. With AlphaFold2 and RoseTTAFold, it is now possible to get reliable predicted structures for abundant sequences.

In this study, we trained a partial-labeled protein design model (BBP) based on ProdCoNN and DenseCPD for protein design from a given structure with a partially labeled sequence. Combining BBP with sequential *Monte Carlo* method, we could sample multiple sequences, and then use AlphaFold2 and RoseTTAFold to predict the structures of the sampled sequences. We compared the predicted structures with the native structure of the wild-type sequence to characterize the foldability of the designed sequences. We used RMSD to measure the similarity between predicted structure to native structure and select a reasonable cutoff value to define designability. Those sequences with predicted structures with RMSD smaller than the cutoff were considered as foldable and used for estimating the designability.

## Methodology and Material

In this paper, we used the sequential *Monte Carlo* method to do sampling based on the prediction probability of each target residue for the protein design part. The previous approaches only use the 3D coordinate of backbone atoms for the sequence prediction without any amino acid information of surrounding residues, which will only output one predicted sequence with one input structure coordinate. However, to estimate the designability, we need to sample multiple sequences from given structure. When sampling a sequence, we used sequential *Monte Carlo* method by sampling one residue at a time. A new residue is sampled conditioning on the residues sampled before it. Thus, we need a model to make predictions of the amino acid type of target residue while the surrounding residue types are partially labeled. The procedure of the sequential *Monte Carlo* method is briefly shown in Fig. 1. In the first step, one single residue is regarded as the target residue, with all others recognized as unknown types. After the target residue is sampled as G(Glycine), the next target residue will be selected, and its type will be predicted based on the result that the previous target residue is known as G. Following this, all the residues along the sequence will be sampled until no unknown type is detected. Since the order of sampling varies and the randomness during the sampling, the diversity of output sequences is guaranteed.

**Figure 1.**
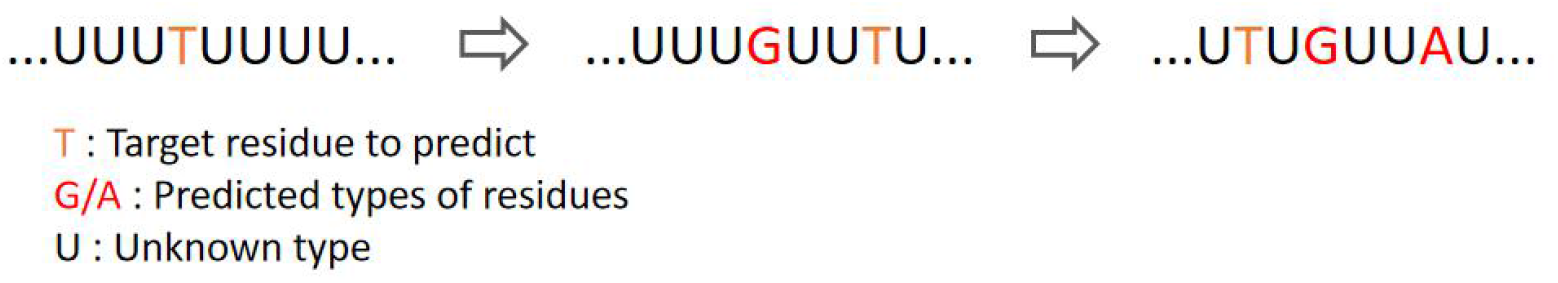
Procedure of protein design using sequential *Monte Carlo*

### Partial-labeled protein design model

We trained a protein design model (BBP) with a partial sequence known to make the prediction based on ProdCoNN[22] and DenseCPD[23]. The data preparation process is similar to the one used in ProdCoNN. The architecture of the neural network applied the DenseNet3D model, which has been shown to improve the model accuracy compared with the simple CNN model in ProdCoNN. We use a gridded box with an edge length of 20 Å centered on the CA atom of the target residue to capture the local structural information around it. The box is gridded with each voxel of size (1 Å × 1 Å × 1 Å). With this resolution, most of the voxels contain no more than one heavy atom. The protein is rotated for each target residue to make the CA-CB bond on the positive z-axis, the CA-N on the x-z plane, and the atom N on the “-x” side. Each atom type is represented by a different channel, like RGB color channels in images. Twenty-seven channels are used, corresponding to atoms C, CA, N, O, OXT, CB atom of target residue, CB atoms of unknown residues, and 20 types of CB atoms of type-known non-target residues. With this setup, the dimension of input data is 20 × 20 × 20 × 27. In this model, some non-target CB atoms in the box are known and labeled as the so-called “partial” here. The number of labeled CB atoms are randomly generated, and the probability of each CB atom being labeled is negatively correlated with the distance from the target residue. In addition, the input data are smoothed using 3D truncated Gaussian functions, so each heavy atom is represented by a Gaussian density function spread over the voxel the atom occupies and the 26 adjacent voxels.

We chose 10,149 protein structures (ID30) with the sequence’s identity lower than 30% from PDB[42]. All the structures were determined by X-ray crystallography with a resolution better than 2.0 Å and did not contain any DNA/RNA/UNK residues. We randomly picked 8135 protein structures and generated 10,000,000 partial-labeled boxes as training data set. 1000 protein structures were chosen, and 1,000,000 partial-labeled boxes were generated as validation data set. 1014 protein structures were chosen, and 200,000 partial-labeled boxes were generated as test data set. Another 50 protein chains (TS50) (252,250 partial-abeled boxes), which are not included in the above data set, are used to test the models and compare with other methods.

The neural network’s architecture is based on the implementation of DenseNet in 3D, which has been proved to be efficient in DenseCPD. The model consists of three Dense blocks with transition blocks between them. In each Dense block, six convolution blocks are concatenated. After batch normalization, ReLU and global average pooling layers are followed. Then, the SoftMax output is generated with 20 numbers representing the probability of being predicted as the 20 types of amino acids. In this paper, we set the learning rate to be 0.0001, the growth rate in the dense block to be 35, and the training batch size to be 50.

### Sampling details

The sampling order is decided based on the calculated entropy along the sequence:

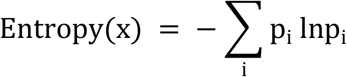

Where P_i_ is the probability that residue x is predicted as amino acid i. And the sampling probability is regulated by

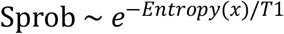

which includes a hyper-parameter temperature T1. The residues with lower entropy have a higher chance to be sampled first, while different sampling iterations could generate different sampling orders.

When the sampling order is decided, the amino acid types will be predicted by the BBP. Here we will not use the output prediction probability from the model directly but modify it with exponential function:

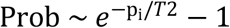

We did this to exaggerate the differences of probabilities, and here another temperature T2 was introduced. We used “minus 1”here to adjust the probability to be zero when the original P_i_ is zero.

To characterize the designability, we calculated the weight of one sampled sequence:

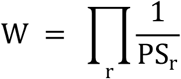

where r goes through all residues, and PS is the probability of amino acid we chose for residue r. In a 100% determined sampling (every residue being predicted as a type of amino acid with 100% probability), PS is always 1, and W is 1. In this case, the size of designability is one. Lower PS values will lead to a larger W and a larger diversity of designability.

We found that some pdb structures could not generate “good”sequences that fold to a similar structure compared with the wild-type sequence structure. We introduced a method to constrain the prediction probability. By analyzing the MSA results of the wild sequence from HHsuite[43], we constrained the prediction probabilities of top 20 residues with high aligned MSA scores. Each residue is constrained to have a 95% chance to be sampled based on the distributions of amino acids from MSA results, and our model predictions still determine the remaining 5%. The problem mentioned above was well solved by this method without reducing the varieties of the sampled sequences a lot.

### Structural prediction of designed sequences

To estimate the foldability of the designed sequences, we used AlphaFold2[40] and RoseTTAFold (pyrosetta version) [41] to predict the 3D structures of the sampled sequences.

Nine pdb structures were selected from the SCOPe database [12], which belong to 7 major classes defined by SCOPe. Respectively, they are two structures from *all alpha proteins*, two structures from *all beta proteins*, one structure from *alpha and beta (a/b), alpha and beta(a+b), multi-domain, membrane and cell surface, coiled coil* and *peptides proteins*. The lengths of the structures range from 76 to 156 residues. All the selected pdb structures are single chains without broken parts in the middle and uncommon amino acids like *selenomethionine*. Also, no binding ligands are included. The details of those structures and their visualizations are shown in Table 1.

**Table 1.**
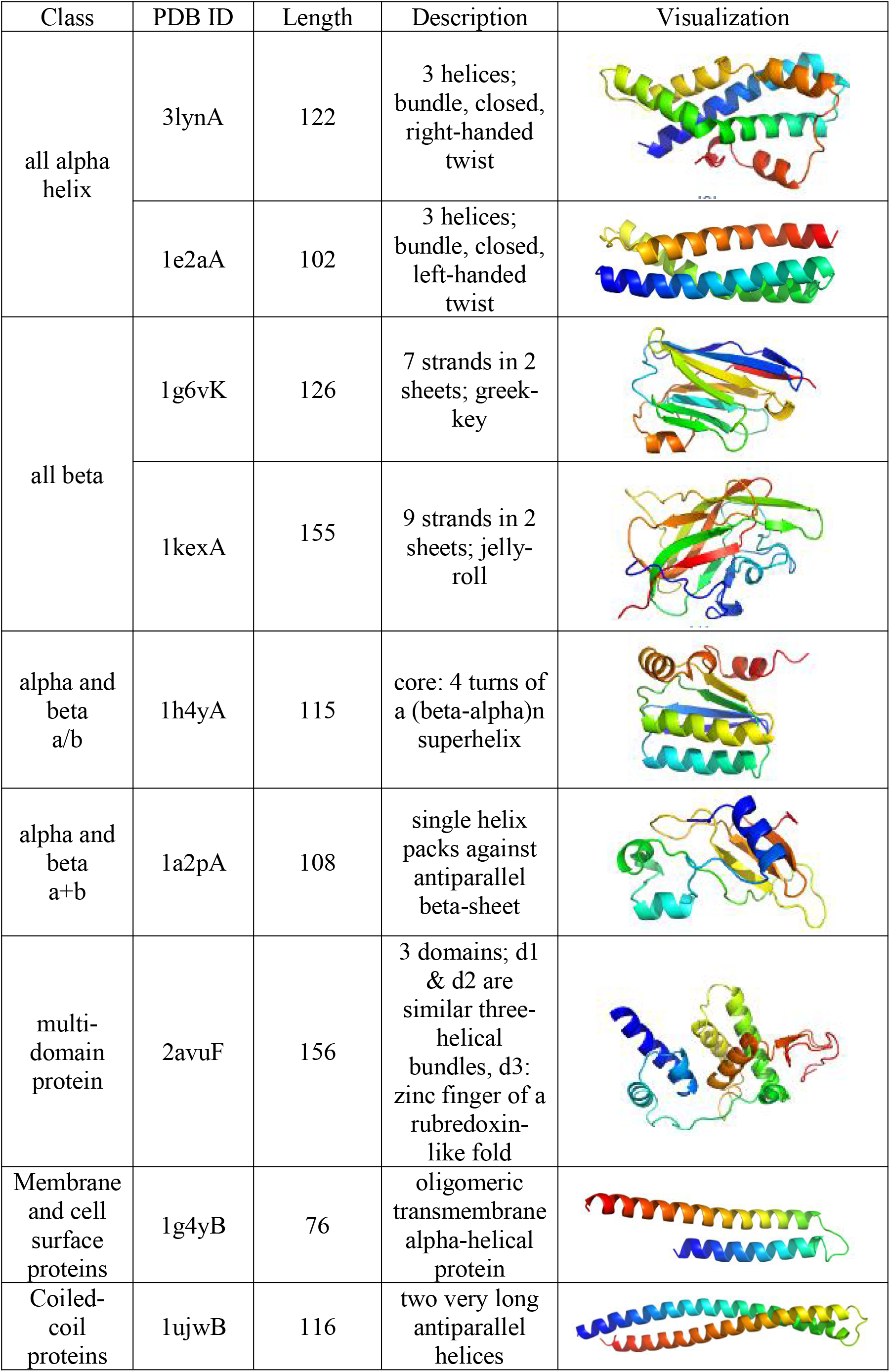
Details of 9 pdb structures selected from SCOPe database

| Class | PDB ID | Length | Description | Visualization |
| --- | --- | --- | --- | --- |
| all alpha helix | 3lynA | 122 | 3 helices; bundle, closed, right-handed twist |  |
|  | 1e2aA | 102 | 3 helices; bundle, closed, left-handed twist |  |
| all beta | 1g6vK | 126 | 7 strands in 2 sheets; greek-key |  |
|  | 1kexA | 155 | 9 strands in 2 sheets; jelly-roll |  |
| alpha and beta a/b | 1h4yA | 115 | core: 4 turns of a (beta-alpha)<sub>n</sub> superhelix |  |
| alpha and beta a+b | 1a2pA | 108 | single helix packs against antiparallel beta-sheet |  |
| multi-domain protein | 2avuF | 156 | 3 domains; d1 & d2 are similar three-helical bundles, d3: zinc finger of a rubredoxin-like fold |  |
| Membrane and cell surface proteins | 1g4yB | 76 | oligomeric transmembrane alpha-helical protein |  |
| Coiled-coil proteins | 1ujwB | 116 | two very long antiparallel helices |  |

## Results and Discussion

### Partial-labeled protein design model (BBP) training result

The BBP model was finished training with 20 epochs. The training loss/accuracy is constantly decreasing/increasing within 20 epochs, while validation loss/accuracy starts to increase/decrease after a few epochs. The minimum validation loss is 1.5326, and the maximum validation accuracy is 0.5182 at the fifth epoch.

We also tested the models of different epochs with the test set and TS50 data set, and the best results were also given at the fifth epoch. We compared the result with another two models we trained: non-labeled model (means no CB atoms are labeled as known) with DenseNet architecture; partial-labeled model but with simple CNN architecture, and the performance is better. This proves that the extra information given by partially labeled CB atoms and DenseNet architecture can both help improve the model. For the following work, we used the model at the fifth epoch to generate sampled sequences.

We also tested the models of different epochs with the test set and TS50 data set, and the best results were also given at the fifth epoch. We compared the result with another two models we trained: non-labeled model (means no CB atoms are labeled as known) with DenseNet architecture; partial-labeled model but with simple CNN architecture, and the performance is better. This proves that the extra information given by partially labeled CB atoms and DenseNet architecture can both help improve the model. For the following work, we used the model at the fifth epoch to generate sampled sequences.

To estimate the designability when we sample the sequences using the *Monte Carlo* method, we tested on five pdb structures(1ahsA, 1eteA, 1i8nA, 2a2lA, 3ejfA). For each structure, we sampled 5000 times independently and used the top 3 predicted probabilities of each residue for sampling. We used the weight of sampled sequences to indicate the designability of the input structure and plot the average weight against sample size for each pdb shown in Fig. 2. The figure shows that the average weight ranges from about 1050 to 10100, indicating a considerable designability for most protein design assignments.

**Figure 2.**
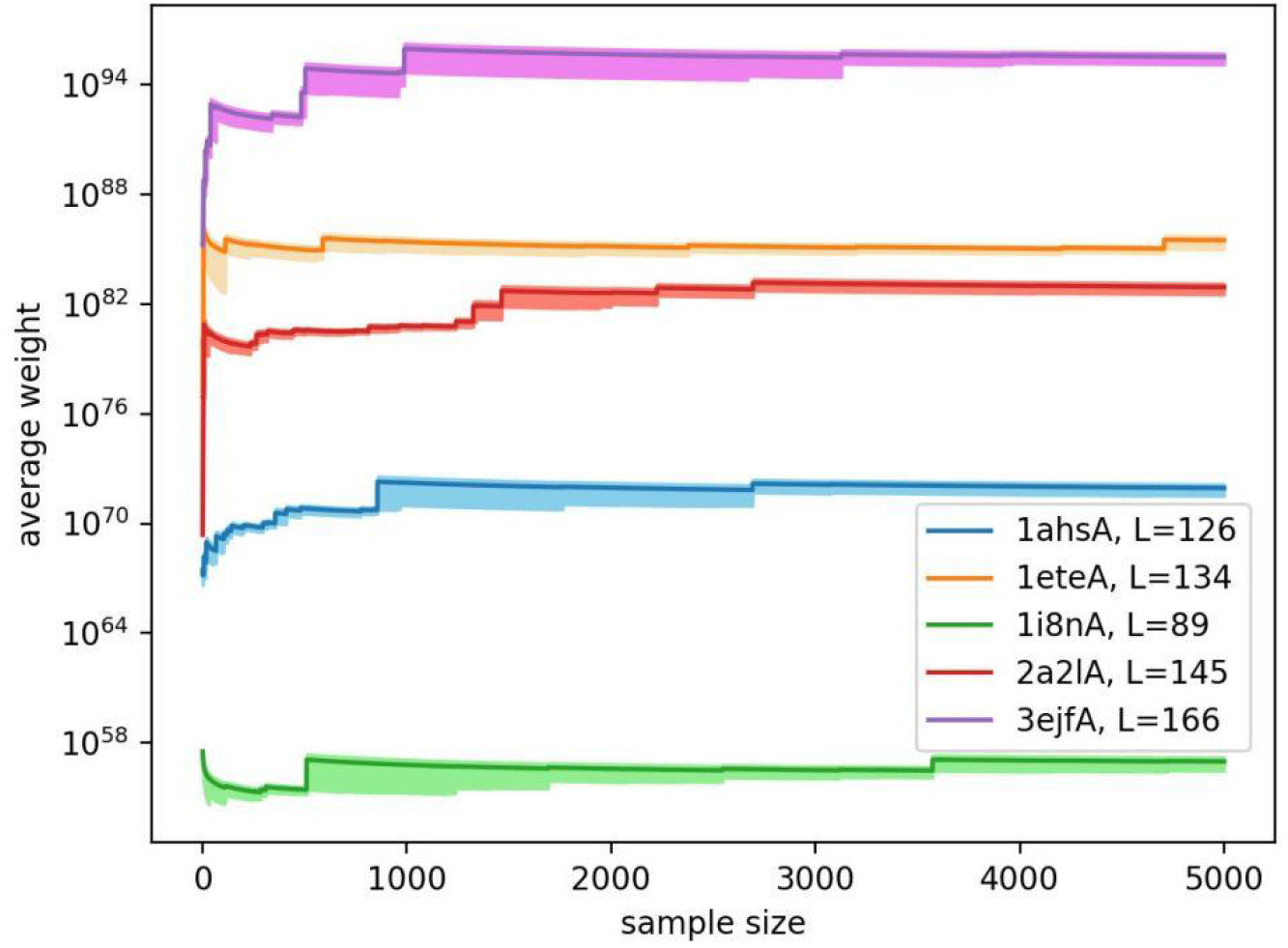
Average weight against sample size for five testing pdb structures, the y-axis is in log scale. The estimated error is shown in lighter color bands, and the lengths of the structures are shown in the legend.

The estimated errors for each pdb are relatively small compared with the absolute value, and longer protein always reasonably lead to a larger designability. Besides, the sampling shows convergence since the weight goes to converged values as sampling proceeds.

### Sequence design and structural prediction

Nine pdb structures selected from SCOPe are used for sampling under two schemes: unconstrained probability and constrained probability. We generated 1000 sampled structures for each pdb structure with the hyper-parameter T1 and T2 set to 1 and 0.05. We analyzed the sequence similarities among the sampled sequences as well as to the wild type sequences. The results are shown in Table 2. For the unconstrained probability, the average sequence similarity to the wild-type sequence for each pdb ranges from 16% to 31%. The average sequence similarity among the sampled sequences ranges from 28% to 36%. For the constrained probability, the correspondent values are range from 24% to 37% and range from 36% to 45%. The constrained probability increases the similarity by around 8%, but they are still below 50% to guarantee the varieties of samplings.

**Table 2.** Sequence similarity of each pdb for unconstrained and constrained designs. SS1: sequence similarity among the sampled sequences; SS2: sequence similarity to the wild sequence

| PDB ID | Unconstrained SS1 | Constrained SS1 | Unconstrained SS2 | Constrained SS2 |
| --- | --- | --- | --- | --- |
| 3lynA | 32% | 37% | 22% | 28% |
| 1e2aA | 33% | 39% | 23% | 30% |
| 1g6vK | 34% | 42% | 23% | 31% |
| 1kexA | 38% | 43% | 31% | 37% |
| 1h4yA | 32% | 38% | 25% | 30% |
| 1a2pA | 36% | 43% | 28% | 36% |
| 2avuF | 30% | 36% | 19% | 24% |
| 1g4yB | 30% | 45% | 16% | 36% |
| 1ujwB | 28% | 36% | 18% | 26% |

For each pdb, we randomly chose 100 out of the 1000 sequences and used AlphaFold2/RoseTTAFold to predict their structures. We defined the predicted structures with real RMSD (native structure of wild-type sequence as reference) lower than 5 Å as foldable and unfoldable vice versa. The percentage of foldable sequences is considered as a foldability score. Table 3 concludes the results of RoseTTAFold and Alphafold, including the real CA RMSD when input is the wild-type sequence, the foldability score, and minimum foldable CA RMSD for both unconstrained and constrained cases.

**Table 3.** Results of prediction for RoseTTAFold (RF) and AlphaFoldF2 (AF)

| PDB ID | RMSD (wild-type seq) |  | Foldability (unconstrained) |  | RMSD<sub>min</sub>(Å) (unconstrained) |  | Foldability (constrained) |  | RMSD<sub>min</sub>(Å) (constrained) |  |
| --- | --- | --- | --- | --- | --- | --- | --- | --- | --- | --- |
|  | RF | AF | RF | AF | RF | AF | RF | AF | RF | AF |
| 3lynA | 0.95 | 0.77 | 14% | 27% | 1.62 | 1.19 | 86% | 87% | 1.16 | 1.06 |
| 1e2aA | 1.08 | 0.79 | 0 | 2% | 6.57 | 5.83 | 72% | 87% | 1.20 | 1.12 |
| 1g6vK | 1.39 | 0.82 | 0 | 0 | 7.52 | 8.26 | 31% | 9% | 1.86 | 3.89 |
| 1kexA | 1.04 | 0.67 | 97% | 40% | 1.26 | 1.65 | 99% | 79% | 1.08 | 1.33 |
| 1h4yA | 1.78 | 1.25 | 16% | 6% | 2.79 | 2.62 | 76% | 44% | 1.58 | 1.70 |
| 1a2pA | 0.72 | 0.29 | 52% | 30% | 1.23 | 1.94 | 98% | 47% | 1.08 | 0.76 |
| 2avuF | 1.55 | 0.81 | 2% | 2% | 4.33 | 3.01 | 65% | 71% | 1.88 | 1.77 |
| 1g4yB | 1.16 | 1.26 | 0 | 3% | 5.45 | 3.66 | 65% | 90% | 1.25 | 0.96 |
| 1ujwB | 1.03 | 1.07 | 0 | 0 | 14.63 | 7.35 | 7% | 2% | 2.49 | 2.09 |

From Table 3, we can conclude that a) For all nine pdb structures, RoseTTAFold and AlphaFoldF2 can work very well to predict the wild sequence to structures with real RMSD lower than 2 Å. b) For sequences designed with unconstrained probability, 4 (2) structures cannot generate any foldable sequences, 1 (3) with a very low foldability of 2-3% by RoseTTAFold (AlphaFoldF2). c) For sequences designed with constrained probability, the foldability of those structures that cannot fold in unconstrained cases increased significantly, like 72% for 1e2aA, 31% for 1g6vK, and 65% for 1g4yB. The only exception is 1ujwB, but we still got improvement from 0 to 7%. d) The foldable minima of RMSD for the constrained case are lower than the unconstrained case, indicating a closer structure to the native. As shown, the unconstrained or constrained probability affects the foldability of designed sequences significantly. We will discuss more details later.

Figs. 3 - 5 show the correlation analysis of several parameters, including weight, predicted and real RMS from RoseTTAFold, predicted tm score (ptm) and real RMS from AlphaFold2. From the observation of RoseTTAFold data of all structures, no apparent correlations between weight and real RMS have been found. A typical example is shown in Fig. 3 for 1a2pA sampled in unconstrained probability. This pdb has evenly distributed foldable and unfoldable cases, and linear regression between the logarithm of weight and real RMS gives zero coefficient indicating no correlation. Fig. 4 shows the correlation between predicted attributes of the two programs and real CA RMS for 5 pdbs. Theoretically, a higher predicted tm score indicates that AlphaFold2 has higher confidence in the prediction. The predicted structure should be a high-quality structure closer to the ground true structure. Here assuming that our designed sequences are well sampled based on the input wild structure coordinates, we can consider the “true” structure of the designed sequence to be the native structure of the wild-type sequence. Therefore, it is well consistent that the predicted ptm is negatively correlated to the real RMS with an R-squared coefficient of 0.6.

**Figure 3.**
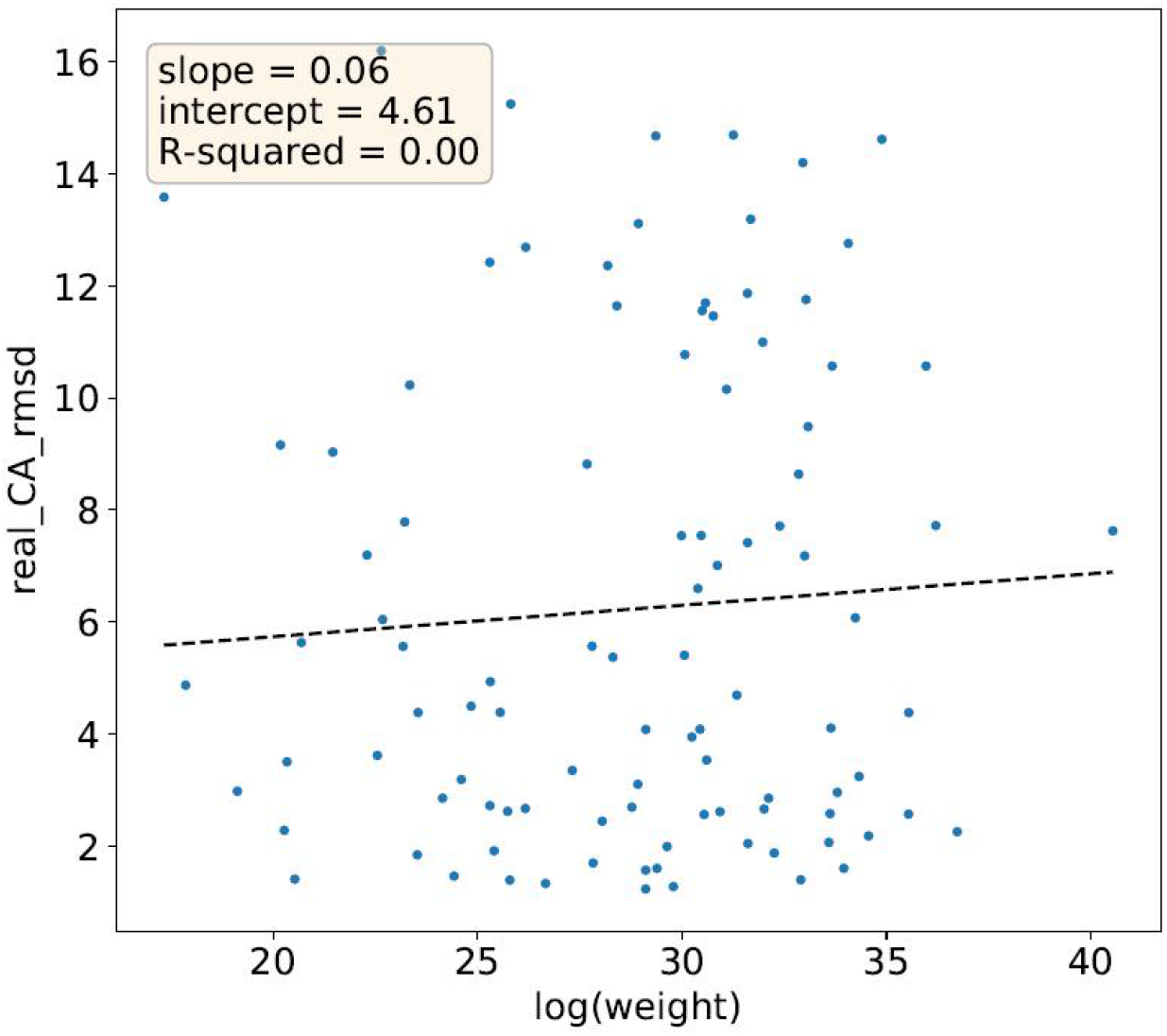
Linear regression of real RMS versus the logarithm of weight for 1a2pA

**Figure 4.**
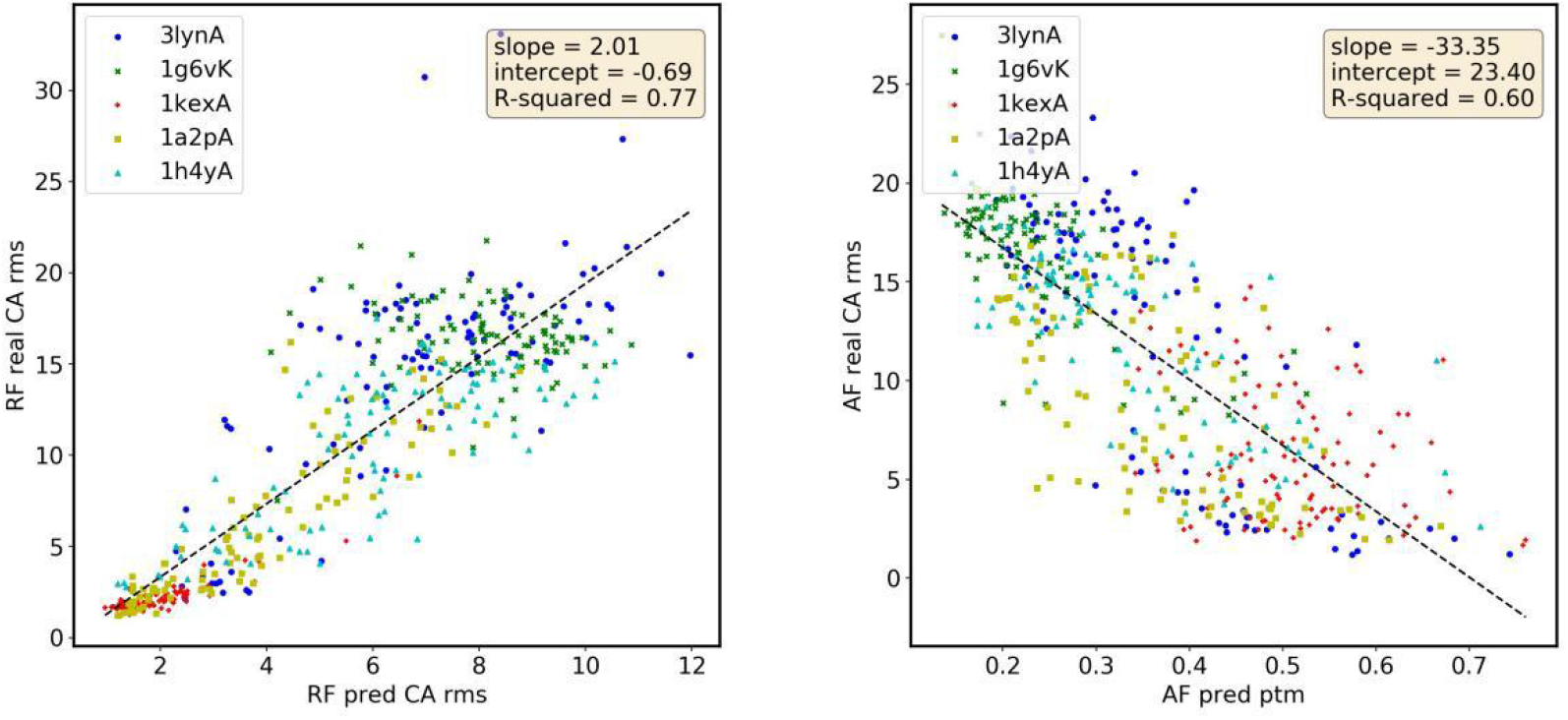
Correlation between predicted attributes of the two programs and real CA RMS for 5 selected structures. Left: RoseTTAFold Right: AlphaFold2. Data are from unconstrained design.

**Figure 5.**
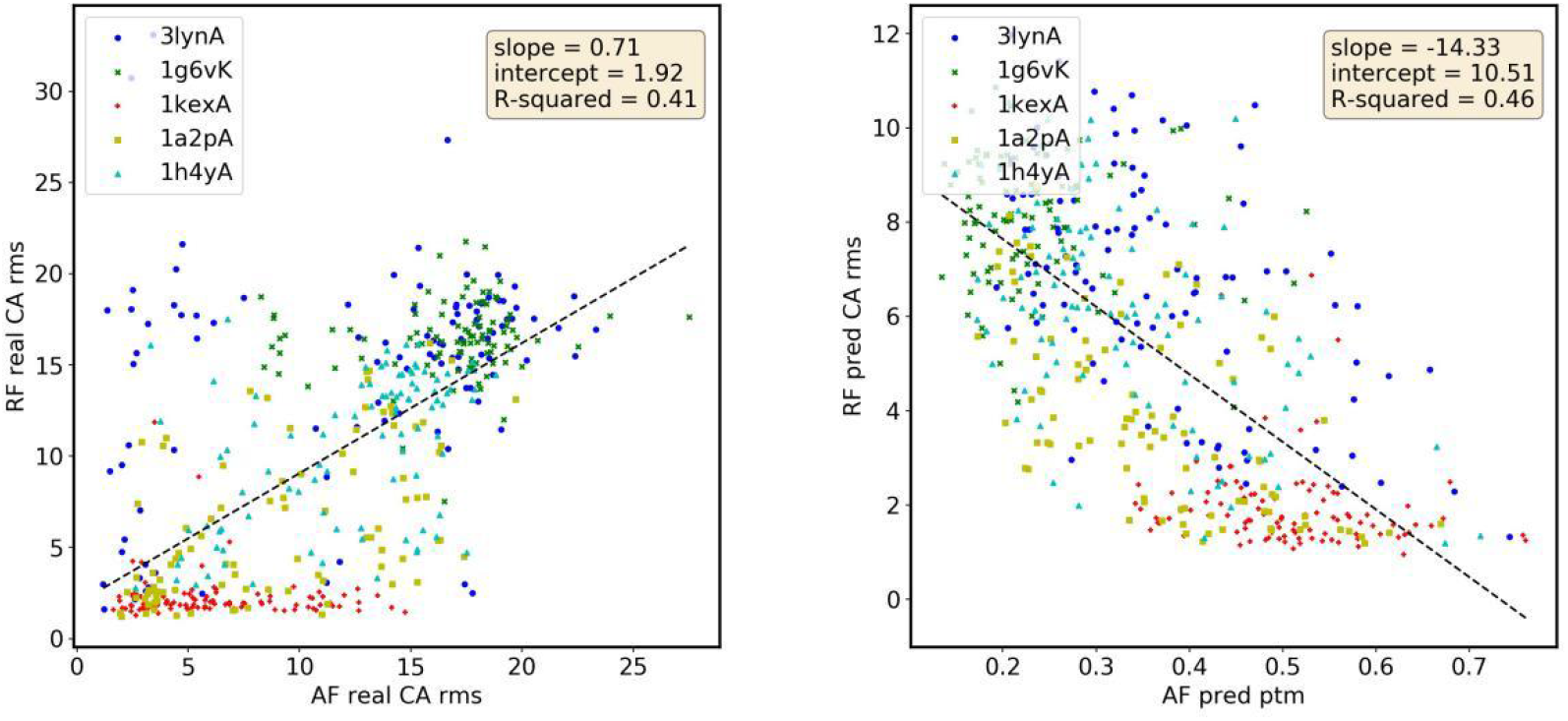
Correlation between the results of the two programs on real RMS(left) and predicted ptm/RMS (right). Data are from unconstrained design.

Similarly, the predicted RMS and real RMS in RoseTTAFold are also correlated with an R-squared coefficient of 0.77. Fig. 5 shows the comparison between the two programs. Real RMS and predicted ptm/RMS give positive correlations, indicating that the two programs are consistent in predictions.

Finally, we discuss how the constrained probability increases the foldability of the designed sequences. We use 3lynA as an example since its foldability increases from 14% (27%) to 86% (87%) by RoseTTAFold (AlphaFold2) in Table 3. In Fig. 6, we selected the wild sequence and 5 designed sequences from constrained sampling, analyzed their MSA results, and plotted the top 10 highly aligned residues for each sequence. The X-axis gives the residue index and amino acid type in each sequence. The Y-axis in the figure shows the frequency that the residue is considered well aligned in MSA results. A higher Y value represents a more conserved residue. The figure shows that the first two designed sequences have very similar frequency patterns with the wild sequence, while they are also well folded with real RMS 1.26 and 2.81. The last three sequences have much lower frequency values, and they are folded with larger RMS or unfolded. The constrained probability can help sample sequences with similar MSA results with the wild-type sequence, especially for the highly conserved residues. And if the highly conserved residues are sampled well or not can significantly determine the foldability of the sequence. In another respect, Fig. 7 plots the logo figure for both unconstrained and constrained foldable sequences. The height of the logos presents the frequencies that each type is sampled. Compared with Fig. 6, we found that most of the highly aligned residue in Fig. 6 also have dominant logos in Fig. 7, like 29(G), 40(L), 52(W), etc. It is not surprising since the highly aligned residues are highly likely to be conserved to certain types, most likely the wild types. On the other hand, the logos in constrained sequences include more dominant logos because of the probability constraint added in sampling. These changes could be of various kinds. Like for 18(D), the frequency is increased in constrained sequences. While for 24(W), the constrained sequences have more chances to be sampled as F or Y, which could be a reasonable adjustment as shown in the two foldable sequences in Fig. 6. However, for 29(H), the foldable unconstrained sequences are never found to be sampled as H, so this could be a redundant restraint. The strategy of adding MSA constraint in sampling can help to sample more foldable sequences, which has been affirmed in Table 3.

**Figure 6.**
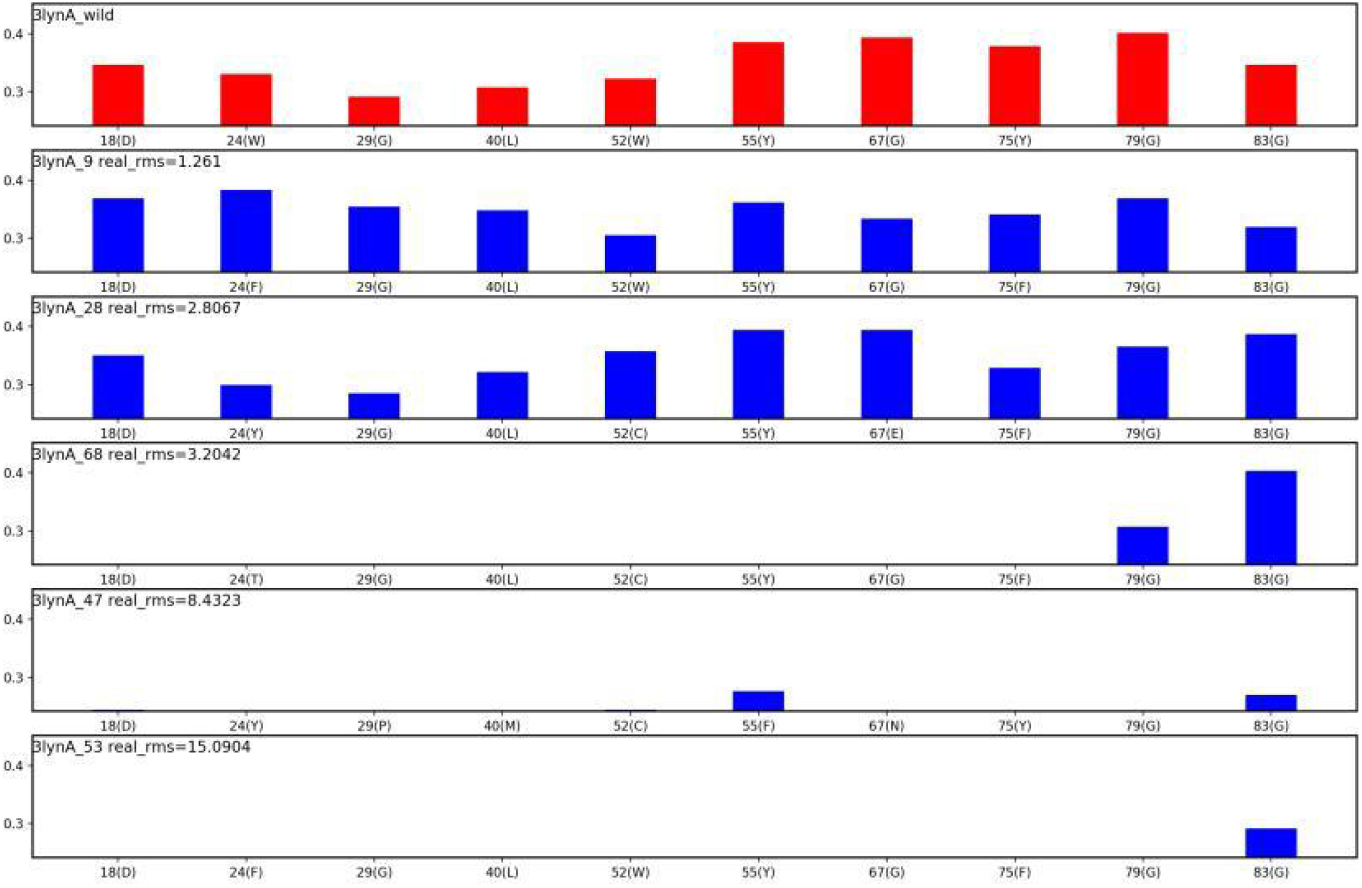
The top 10 of the highly aligned residue from MSA results for wild sequence and five constrained designed sequences

**Figure 7.**
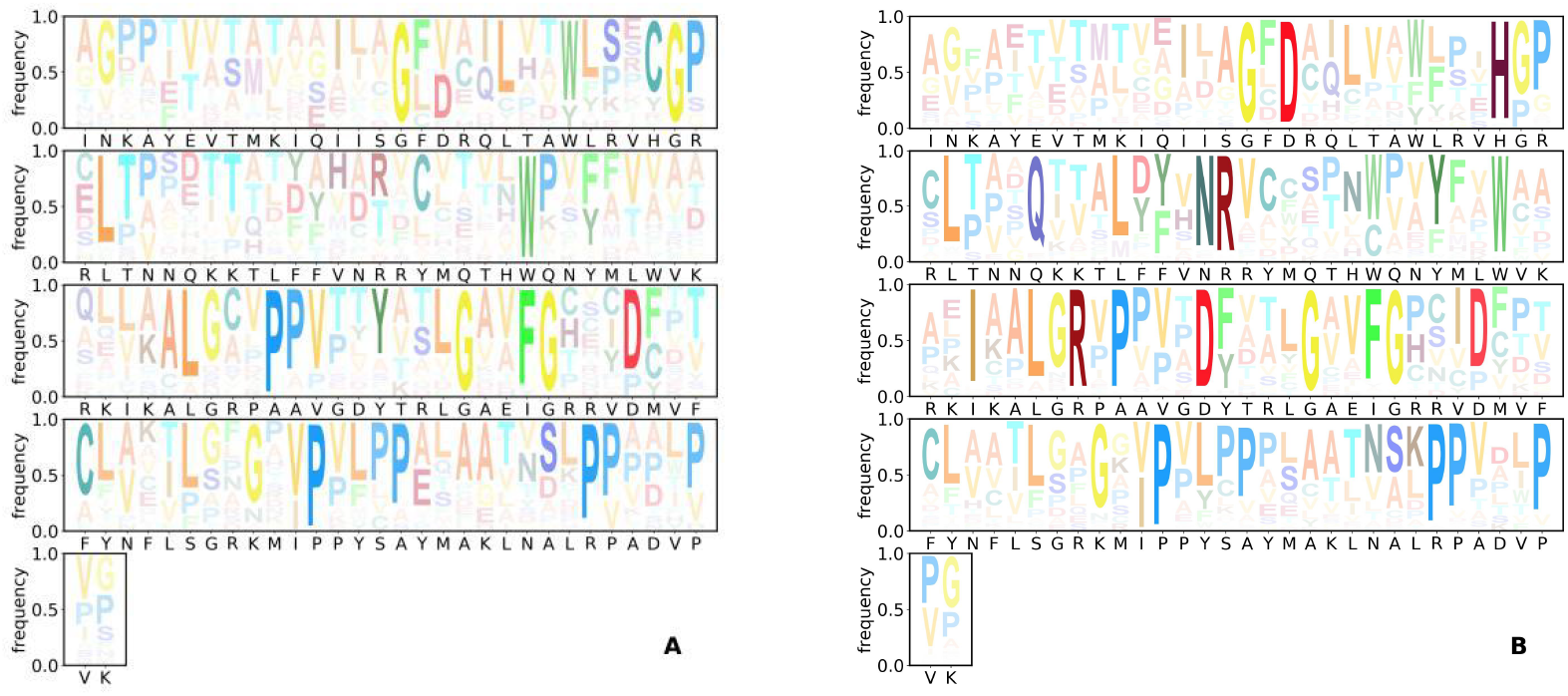
Logo plots indicating the sampling frequencies of each residue type for unconstrained design(A) and constrained design(B)

## Conclusion

With the trained partial-labeled protein design model, we sampled designed sequences of nine pdb structures from seven protein families. Then we used AlphaFold2 and RoseTTAFold programs to estimate the foldability of the designed sequences. Both the sequence similarities among the sampled sequence and relative to the wild-type sequence are below 40%, which guarantees the diversity and variability of the sequences distributed in the designable space. The resultant weights representing the magnitude of designability vary dramatically between different proteins, indicating the designability variation of these proteins. From the structure prediction results, foldable sequences were found in seven proteins for AlphaFold2 and five proteins in RoseTTAFold, with highest foldable percentage as 97%. After modifying the sequence prediction probability based on the MSA results of wild-type sequence, foldable sequences were found in all nine proteins for both AlphaFold2 and RoseTTAFold. The comparison of MSA results between wild-type and sampled sequences shows that the corrected probability of possible critic residues improved foldability significantly.

